# Alignment-free machine learning approaches for the lethality prediction of potential novel human-adapted coronavirus using genomic nucleotide

**DOI:** 10.1101/2020.07.15.176933

**Authors:** Rui Yin, Zihan Luo, Chee Keong Kwoh

## Abstract

A newly emerging novel coronavirus appeared and rapidly spread worldwide and World Health Organization declared a pandemic on March 11, 2020. The roles and characteristics of coronavirus have captured much attention due to its power of causing a wide variety of infectious diseases, from mild to severe on humans. The detection of the lethality of human coronavirus is key to estimate the viral toxicity and provide perspective for treatment. We developed alignment-free machine learning approaches for an ultra-fast and highly accurate prediction of the lethality of potential human-adapted coronavirus using genomic nucleotide. We performed extensive experiments through six different feature transformation and machine learning algorithms in combination with digital signal processing to infer the lethality of possible future novel coronaviruses using previous existing strains. The results tested on SARS-CoV, MERS-Cov and SARS-CoV-2 datasets show an average 96.7% prediction accuracy. We also provide preliminary analysis validating the effectiveness of our models through other human coronaviruses. Our study achieves high levels of prediction performance based on raw RNA sequences alone without genome annotations and specialized biological knowledge. The results demonstrate that, for any novel human coronavirus strains, this alignment-free machine learning-based approach can offer a reliable real-time estimation for its viral lethality.

## Introduction

Coronaviruses are positive, single-stranded RNA virus and have been identified in humans and animals. They are categorized into four genera: *α, β, γ* and *θ* [1]. Previous phylogenetic analysis revealed a complex evolutionary history of coronavirus, suggesting ancient origins and crossover events that can lead to cross-species infections [2] [3]. Bats and birds are a natural reservoir for coronavirus gene pool [4]. The mutation and recombination play critical roles that enable cross-species transmission into other mammals and humans [5]. Human coronavirus (HCoV) was first identified in the mid-1960s [6], and up to now, seven types of coronavirus can infect people. Four of them, i.e., HCoV-229E, HCoV-NL63, HCoV-OC43 and HCoV-HKU1, usually cause mild to moderate upper-respiratory tract illness like common cold when infecting humans [7]. The other three members include severe acute respiratory syndrome coronavirus (SARS-CoV) and middle east respiratory syndrome coronavirus (MERS-CoV) and the most lately severe acute respiratory syndrome coronavirus 2 (SARS-CoV-2), also known as COVID-19 coronavirus. They all belong to Betacoronavirus that led to the epidemics or pandemics [8] [9] [10].

Emerging in November 2002 in Guangdong province, China, SARS caused 8,096 human infections with 774 deaths by July 2003 [11]. MERS was first reported in Saudi Arabia in September 2012 and finally resulted in 2,494 human infections by November 2019 [12]. Recently, a novel coronavirus named COVID-19 is emerging and spreading to 215 countries or territories on June 12, 2020, leading to 7,390,702 confirmed cases with 417,731 deaths according to the World Health Organization [13]. Though precautions such as lockdown of cities and social distance have been taken to curb the transmission of COVID-19, it spreads far more quickly than the SARS-CoV and MERS-CoV diseases [4] [14]. To make matters worse, the number of infected cases still increases rapidly and the global inflection point about COVID-19 is unknown. Like other RNA viruses, e.g. influenza virus [15], coronaviruses possess high mutation and gene recombination rates [16], which makes constant evolution of this virus with the emergence of new variants. From SARS in 2002 to COVID-19 in 2019, coronaviruses have caused high morbidity and mortality, and unfortunately, the fast and untraceable virus mutations take the lives of people before the immune system can produce the inhibitory antibody [17]. Currently, no miracle drug or vaccines are available to treat or prevent the humans infected by coronaviruses [18] [19]. Therefore, there is a desperate need for developing approaches to detect the lethality of coronaviruses not only for COVID-19 but also the potential new variants and species. This would facilitate the diagnosis of coronavirus clinical severity and provide decision-making support.

The detection of viral lethality has already been explored in influenza viruses [20]. Through a meta-analysis of predicting the virulence and antigenicity of influenza viruses, we can infer the lethality of the virus timely to improve the current influenza surveillance system [21]. Regarding the risk of novel emerging coronavirus strains, much attention has been captured to investigate the lethality or clinical severity of new emerging coronavirus. Typically, epidemiological models are certainly built to estimate the lethality and the extent of undetected infections associated with the new coronaviruses. Bastolla suggested an orthogonal approach based on a minimum number of parameters robustly fitted from the cumulative data easily accessible for all countries at the John Hopkins University database to extrapolate the death rate [22]. Bello-Chavolla et al. proposed a clinical score to evaluate the risk for complications and lethality attributable to COVID-19 regarding the effect of obesity and diabetes in Mexico [23]. The results provided a tool for quick determination of susceptibility patients in a first contact scenario. Want et al. leveraged patient data in real-time and devise a patient information based algorithm to estimate and predict the death rate caused by COVID-19 for the near future [24]. Aiewsakun et al. performed a genome-wide association study on the genomes of COVID-19 to identify genetic variations that might be associated with the COVID-19 severity [25]. Moreover, Jiang et al. established an artificial intelligence framework for data-driven prediction of coronavirus clinical severity [26]. The development of computational and physics-based approaches has relieved the labors of experiments by utilizing epidemiological and biological data to construct the model. However, direct evaluation of potential novel coronavirus strains for their lethality is crucial when clinicians are forced to make difficult decisions without past specific experience to guide clinical acumen. Inferring the lethality of novel coronavirus is possible by identifying the patterns from a large number of coronavirus sequences.

In this paper, we propose alignment-free machine learning-based approaches to infer the lethality of potential novel human-adapted coronavirus using genomic sequences. The main contribution is that we formulate the problem of estimating the lethality of human-adapted coronavirus through machine learning approaches. By leveraging some appropriate feature transformation, we can encode genomic nucleotides into numbers that allow us to convert it into a prediction task. The experimental results suggest our models deliver accurate prediction of lethality without prior biological knowledge. We also performed phylogenetic analysis validating the effectiveness of our models through other human coronaviruses.

## Materials and Methods

### Problem formulation

The pandemic of novel coronavirus COVID-19 has caused thousands of fatalities, making tremendous treats to public health worldwide. The society is deeply concerned about its spread and evolution with the emergence of any potential new variants, that would increase the lethality. Typically, lethality refers to the capability of causing death. It is usually estimated as the cumulative number of deaths divided by the total number of confirmed cases. Among all the human-adapted coronaviruses, MERS-CoV caused the highest fatality rate of 37% [12], followed by the SARS-CoV with 9.3% fatality rate [27]. In comparison, COVID-19 indicates a lower mortality rate of 5.5% [13]. The lethality rate of COVID-19 is likely to decrease with better treatment and precautions. In this paper, we mainly focus on these three types of human-adapted coronavirus and define the degree of viral lethality in terms of historical fatality rates. As a result, MERS-CoV strains are high lethal while SARS-CoV and COVID-19 strains are middle and low lethal, respectively.

### Data collection and preprocessing

Genomic nucleotide sequences of three different coronaviruses with the human host are downloaded from National Center for Biotechnology Information on April 30, 2020 [28]. Duplicate sequences and incomplete genomes with a length smaller than 20000 are removed from the collection to address the possible issues raised from sequence length bias. Some SARS-CoV strains from the laboratory are included that are cultivated in Vero cell cultures to enrich the training samples. Finally, we end up with 321, 351, 1638 samples for MERS-CoV, SARS-CoV and SARS-CoV-2. In addition, we also collect the genomic data of other four human coronaviruses with 27, 64, 32 and 142 strains for HCoV-HKU1, HCoV-NL63, HCoV-229E and HCoV-OC43, respectively. Apart from the four symbolic bases (A, C, T, G) of each strain, we have degenerate base symbols that are an IUPAC representation [29] for a position on genomic sequences, which could denote multiple possible alternatives. These degenerate base symbols contain W (A, T), S (C, G), M (A, C), K (G, T), R (A, G), Y (C, T), B (C, G, T), D (A, G, T), H (A, C, T), V (A, C, G), N(A, C, T, G), where the letters in the bracket are alternative nucleotide representing degenerate bases. We randomly substitute these degenerate bases so that genomic sequences can be mapped into discrete numerical representation through feature transformation.

### Feature transformation

Numerical representations have been successfully employed in the field of bioinformatics [30] [31], mapping biological sequences into real-value vector space that the information or pattern characteristic of the sequence is kept in order. This is important as the existing machine learning approaches can only deal with vectors but not sequence samples. Several methods are proposed that convert genomic sequences into numerical vectors, e.g., the fixed mapping between nucleotides and real numbers without biological significance [32], based on physio-chemical properties [33], deduction from doublets or codons [34], and chaos game representation [35]. To accommodate comprehensive analysis and comparison, we adapt different types of numerical representations for biological RNA sequences. Randhawa et al. [36] showed that “Real”, “Just-A” and “Purine/Pyrimidine (PP)” numerical representation yield better performance over other methods for DNA classification, which are included for analyzing genomic data. The electro-ion interaction potential (EIIP) and nearest-neighbor based doublet are incorporated that are based on physio-chemical properties and nearest-neighbor values respectively. Apart from the aforementioned one-dimensional representation, we have introduced 2D Chaos Game Representation (CGR) into feature transformation of original sequences.

The real number representation is a fixed transformation technique that we obtain values of four bases as: adenine (A) = −1.5, thymine (T) = 1.5, cytosine (C) = 0.5, and guanine (G) = −0.5 [37]. It is efficient in finding a complementary strand of DNA/RNA sequence and can endure complementary property. Just-A” method maps the four bases into binary classification as the presence of adenine is labeled 1, while others are 0 [38]. PP representation is a DNA-Walk model that shows nucleotides sequences in which a step is taken upwards if the nucleotide is pyrimidine with T/C = 1, or downward if it is purine with A/G = −1 [39]. EIIP describes the distribution of the energy of free elections along with nucleotide sequences that a single EIIP indicator sequence is formed through replacing its nucleotides, where A=0.1260, C=0.1340, G=0.0806, and T=0.1335 [40]. The sequence-to-signal mapping for nearest-neighbor based doublet representation is illustrated in [34], where the last position is followed by the first in the sequence. Lastly, CGR is a method proposed by Jeffrey [41] that has been successfully used for a visual representation of genome sequence patterns and taxonomic classification [42] [43]. The CGR images of RNA/DNA sequences are drawn in a unit square. The four vertices of the square are labeled by four nucleotides. The first nucleotide of the sequence is plotted halfway between the center of the square and the vertex representing this nucleotide. The next base is mapped into the image that the coordinate is assigned halfway between the previous point and the vertex corresponding to the previous nucleotide. The mathematical formulation of the successive points that calculates the coordinates in the CGR of the sequences is described below:

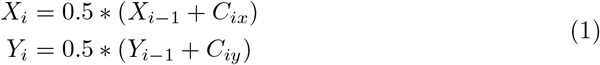

where *C_ix_* and *C_iy_* denote the *X* and *Y* coordinates of the vertices matching the nucleotide at position *i* of the sequence, respectively.

### Model construction

Machine learning has been utilized in many aspects of viral genomic analysis, e.g., antigenicity prediction of viruses [21], genome classification of novel pathogens [43], reassortment detection [44], receptor binding analysis [45] and vaccine recommendation [46], etc. With increasingly available genomic sequences, it will play more critical roles in helping biologists to analyze large, complex biological data for prediction and discovery. In this work, we provide a comprehensive analysis of the lethality inference of potential human-adapted coronaviruses via alignment-free machine learning approaches. The proposed methods not only contain traditional machine learning models but also deep learning techniques in combination with Discrete Fourier Transform for genome analysis. Six different types of numerical representations are implemented in comparison with the predictive performance of machine learning models. Traditional machine learning models consist of logistic regression (LR), random forest (RF), K-nearest neighbor (KNN) and neural network (NN) [47], while three variants of convolutional neural network (CNN) and two types of recurrent neural network (RNN) are leveraged. The CNN models contain AlexNet [48], VGG [49] and ResNet [50].

Following the choices of five one-dimensional numerical representation for viral sequences, digital signal processing is introduced through DFT techniques. We assume that the number of input sequence is *n* and all the sequences have the same length *l*. For each sequence *S_i_* = (*S_i_*(0), *S_i_*(2),…, *S_i_*(*l* – 1)), where 1 ≤ *i* ≤ *n, S_i_*(*k*) ∈ {A, C, T, G} and 0 ≤ *k* ≤ *l* – 1, the corresponding discrete numerical representation is formulated as

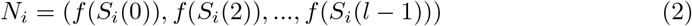

where *f* (*S_i_*(*k*)) denotes the numerical value after mapping by function *f* (·) at the position k of nucleotide sequence *S_i_*. The signal *N_i_* computed after DFT is represented as vector *F_i_*. The formulation of *F_i_* is presented below. We define that the magnitude vector that corresponds to the signal *N_i_* as *M_i_*, where *M_i_* is the absolute value of *F_i_*.

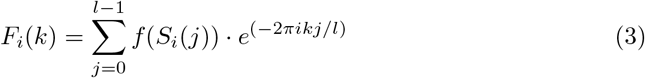

Typically, the length of numerical digital signal *N_i_* is equal to the magnitude spectrum *M_i_* that is originated from the length of the genomic sequence. However, the input genome sequences are in different lengths, thus they need to be length-normalized after DFT. Median length-normalization is leveraged for the input digital signals using zero padding. We employ anti-symmetric padding that begins from the last position if the input sequences are shorter than the median length, these short signals are extended to the median length with zero-padding, while the longer sequences are truncated after the median length.

As for the two-dimensional numerical representation, i.e., CGR, a point that corresponds to a sequence of length l will be contained within a square with a side of length 2^-*l*^. We assume a square CGR image is generated with a size of 2^*k*^ ×2^*k*^ matrix, where *k* is the parameter that determines the size of the image. The frequency of occurrence of any oligomer in a sequence can be obtained by partitioning the CGR space into small squares. Therefore, the number of CGR points in each unit square of 2^*k*^ × 2^*k*^ grid is equal to the number of occurrences of all possible *k*-mers in the sequence. By counting the frequency of CGR points, it is possible to calculate oligonucleotide frequencies at various grid resolutions. We define the element *a_j_* as the number of points that are located in the corresponding sub-square *j*, where 1 ≤ *j* ≤ 2^2*k*^. Each sequence will be mapped into a 2^*k*^ × 2^*k*^ dimensional vector space based on CGR.

### Implementation and evaluation

We implement all the models by Scikit-learn [51] and PyTorch [52]. We utilize the retrospective way to train and test the model since the isolation time of viral strains are available. For each type of coronavirus, the samples generated from strains isolated before the year *N* are used to build the model, while those generated after the year *N* are for testing. The year *N* is determined based on the condition that divides training and testing set in a rough 0.8:0.2 ratio. The 5-fold cross-validation is performed in the training process and the independent testing set is used for validation of our models. This test can truly reflect the ability of the models in applications to predict viral lethality for future strains. The parameters are set by default with traditional machine learning models. For deep learning-based models, we apply stochastic gradient descent with a minimum batch size of 64 for optimization. The drop-out (rate = 0.9) strategy is carried out with a 0.001 learning rate and all the models are fit for 50 training epochs. The predictive performance is evaluated by accuracy, precision, sensitivity, and F1 score of all models in the prediction tasks of coronavirus lethality.

## Results

### Genome composition of SARS-CoV, MERS-CoV and SARS-CoV-2

We first analyzed the composition of the RNA genome of the three human-adapted coronaviruses. Fig 1 portrays the average distribution and variance of the nucleotides. We can observe that the proportion of A and T (in replacement of U) is high, while C and G are relatively low for all human coronaviruses. Interestingly, it is suggested that the high T and low C proportions of human coronaviruses are quite variable and act like communicating vessels. T goes up when C decreases and vice versa. The composition of T ranges from 0.139 to 0.552 while C makes the opposite movement from 0.374 to 0.107 respectively among all human coronavirus. If we look into individual types, the SARS-CoV-2 as a novel human pathogen follows some typical composition of nucleotides but it is also characterized by some differences. We find that SARS-CoV-2 presents a higher variance compared with MERS-CoV and SARS-CoV. This is probably the rapid and widespread transmission of SARS-CoV-2 accelerates its evolution when infected with humans. More strains are generated differently from their ancestor clade. However, it is more pronounced of the nucleotide bias in the unpaired regions of the structured RNA genome, which may indicate a certain biological function of these special sequence signatures. Some studies have revealed that a clear difference in the magnitude of the nucleotide bias of the coronavirus genomes is likely to relate to the mechanism of subgenomic mRNA synthesis and the exposure of single-stranded RNA domains [53] [54]. The evidence shows that cytosine discrimination and deamination against CpG dinucleotides are the driving force that outlines the coronaviruses over evolutionary times [55]. It is indicated that the atypical nucleotide bias could reflect distinct biological functions that are the direct cause of the characteristic codon usage in these viruses [56]. Therefore, the analysis of the nucleotide and codon usage in coronaviruses can not only exhibits the clues on potential viral evolution but also improves the understanding of the viral regulation and promotes vaccine design.

**Figure 1.**
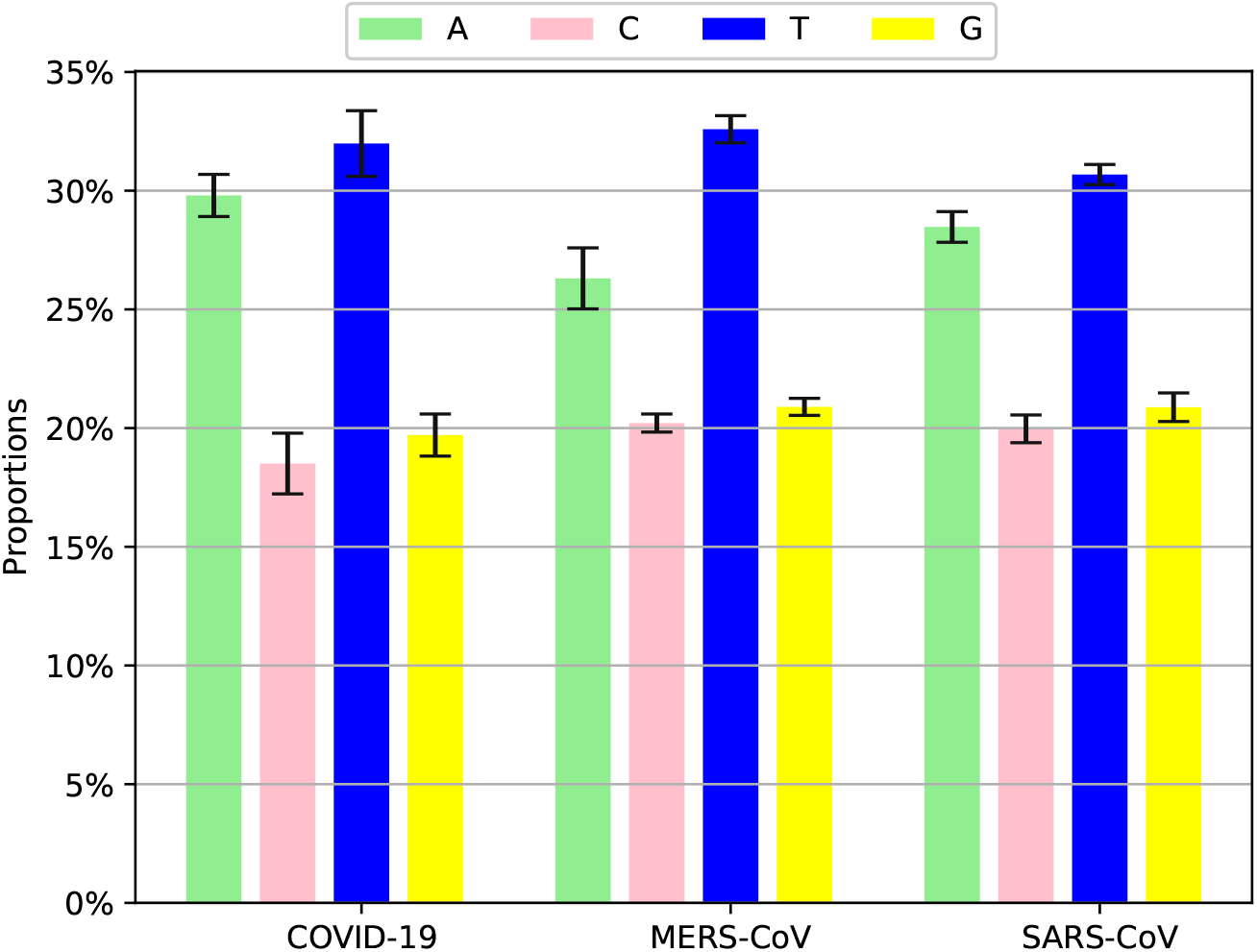
Composition of nucleotide sequence for SARS-CoV, MERS-CoV and SARS-CoV-2.

### K-mer-based classifier study

Studies have investigated the role of *k*-mer frequency for fast and accurate classification of viral genomes [57]. We experiment values for *k*-mer of length *k* =1, 2, …, 7 on different classifiers to measure the prediction accuracy. To explore the *k*-mer frequency patterns in distinct coronaviruses, we curate independent testing set to assess the performance. Fig 2 shows the predictive accuracy across seven machine learning algorithms, at different values of *k*. Fig 2 portrays the performance curves by deep learning models (in the left), and the results via traditional machine learning models (in the right). Overall, our proposed methods obtain an average accuracy of 0.956. However, we can observe the traditional machine learning methods exceed 0.98 in accuracy at all *k* values, whereas there is a different story for deep learning models. It is shown that VGG-19 achieves the best results, while the accuracy could be as low as 0.8 using ResNet-34 when *k* = 2. We can conclude that for these data the traditional machine learning methods outperform deep learning models almost at all levels of *k* with less fluctuation. Considering the tradeoff between computational cost and performance, the k-mer value 6 is used for the results of experiments with CGR representation.

**Figure 2.**
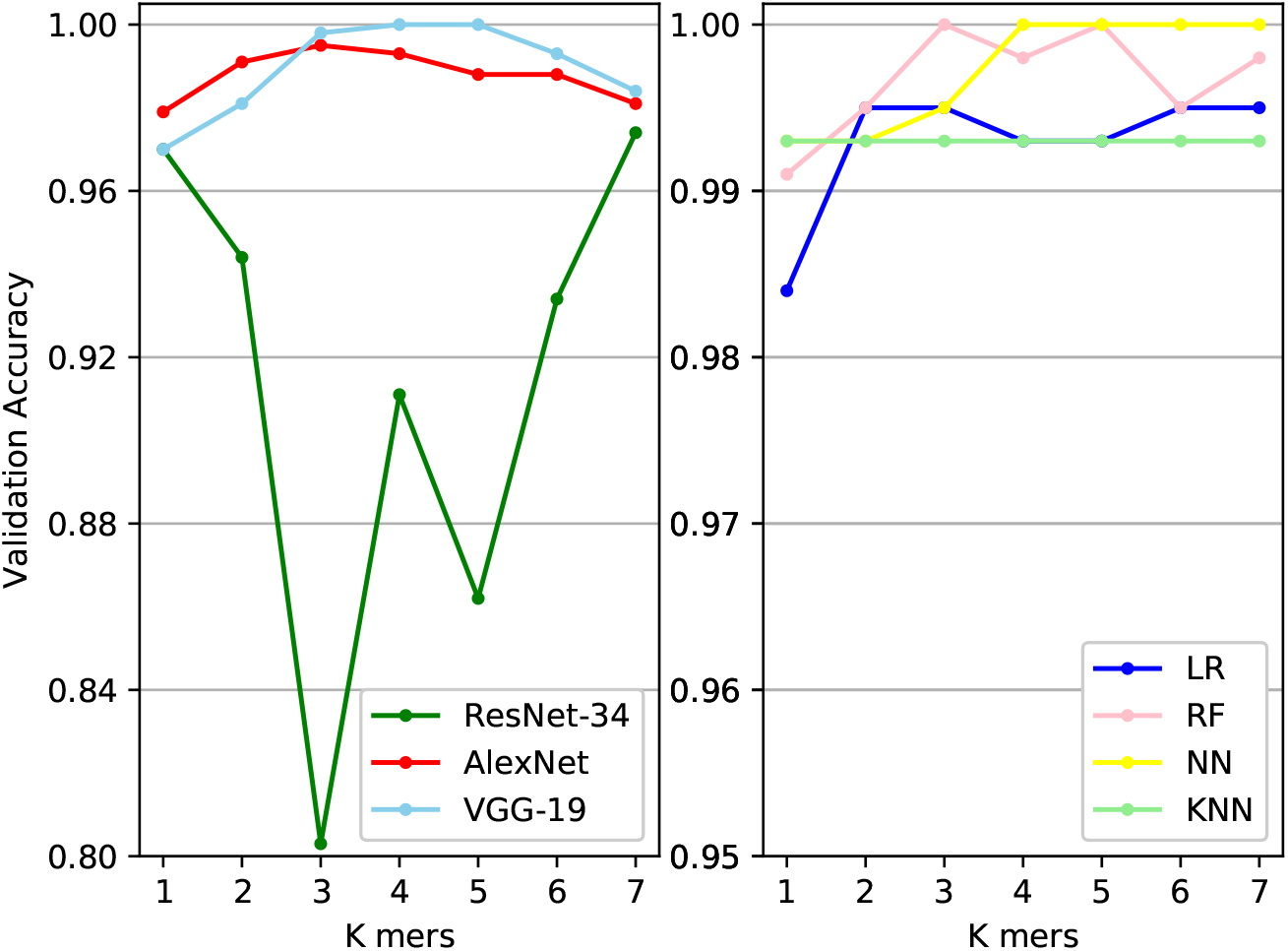
The prediction accuracy across seven machine learning classifiers at different values of *k*.

### Comparative performance

We analyzed the effect on viral lethality prediction via different numerical representations for RNA sequences using machine learning approaches. The dataset used is the same as those in Fig 2. The results along with the average scores for all numerical representations and classifiers are summarized in Table 1. As can be observed from Table 1, for all numerical representations, the average scores are high over all measures. The best performance is achieved when using CGR representation, which yields an average accuracy of 0.985 in the testing set. Surprisingly, we can obtain an average accuracy of 0.967 even with a single nucleotide numerical representation “Just-A”. At the individual classifier level, traditional machine learning methods display an apparent advantage over deep learning models. Logistic regression and neural network can achieve 100% accuracy for all numerical representations, whereas the prediction accuracy ranges from 0.679 to 0.993 implemented by Resnet34, VGG19 and AlexNet. At this point, this is probably because deep learning algorithms need a large amount of data to understand the pattern. In addition to performing higher accuracy, machine learning models are computationally cheaper in this task, e.g. in CGR representation, it takes much longer time for deep learning models (ResNet:4min41s, AlexNet:1min21s, VGG:55s) than classical machine learning methods on average (KNN:1min56s, NN:1min30s, LR:1min7s, RF:12s). Overall, our results suggest that all these numerical representations are effective for modeling to differentiate the degree of the lethality of human coronavirus.

**Table 1.**
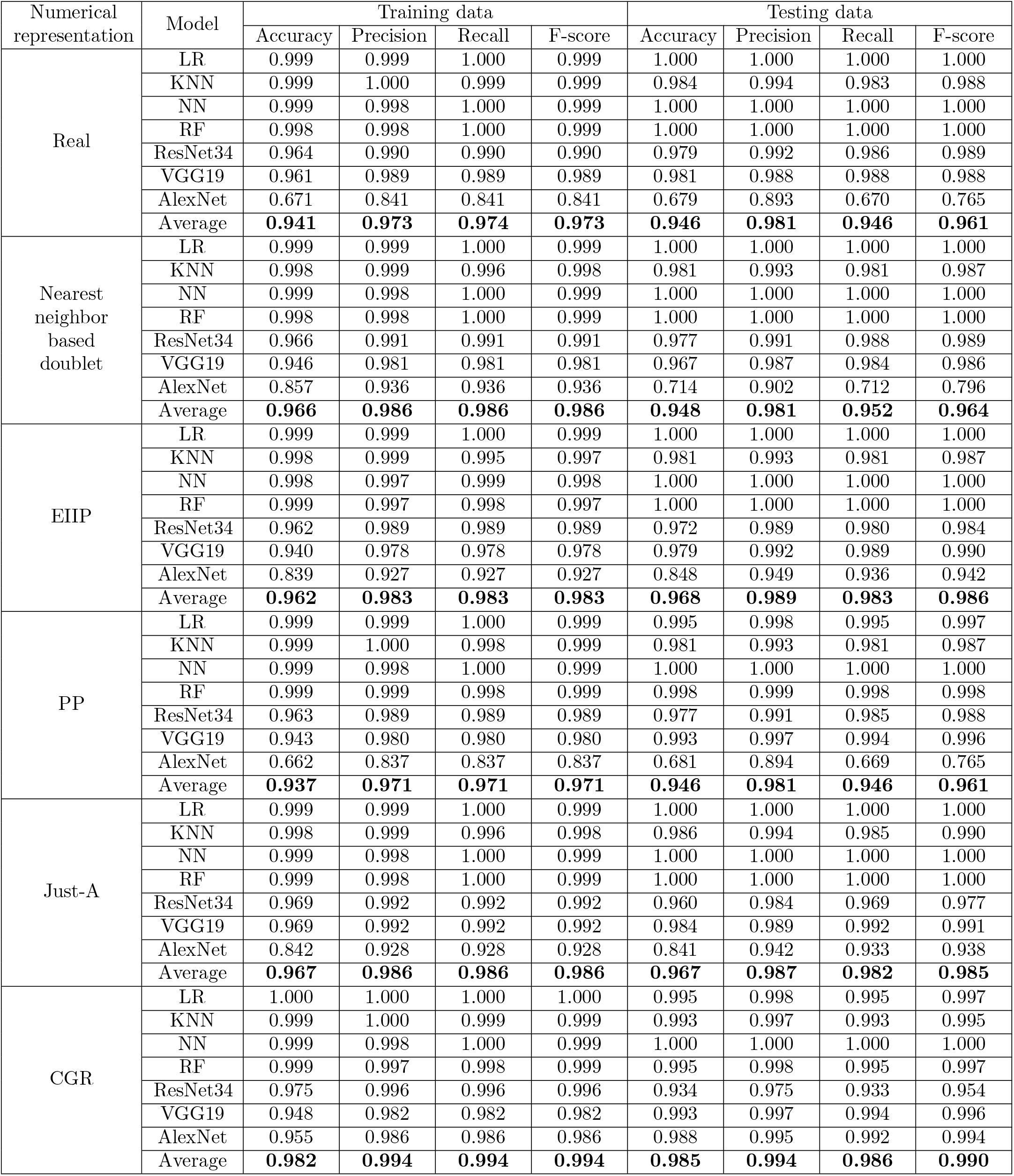
The comparative performance for the lethality prediction of human-adapated coronaviruses via seven different classifierss. Average results for each numerical representation are in bold.

### Validation on other human coronaviruses

We test the ability of our models to identify the lethality of other different human coronaviruses, i.e., HCoV-229E, HCoV-NL63, HCoV-OC43, and HCoV-HKU1. The training process is implemented on the former three types of coronavirus data. For every test dataset, we use CGR as the numerical representation with all classifiers to predict the lethality. Interestingly, the results show that, on average, 28 out of 32 HCoV-229E, 59 out of 64 HCoV-NL63, 134 out of 142 HCoV-OC43 and 25 out of 27 HCoV-HKU1 strains identified have closer lethality with SARS-CoV-2, while the rest strains are labeled middle or high. This suggests that, overall, other test human coronaviruses have lower severity than MERS-CoV and SARS-CoV. Evidence has revealed that HCoV-OC43 and HCoV-HKU1 are associated with mild to moderate upper respiratory tract illness with about 0.1% fatality [58]. These infections may be asymptomatic and are considered the second common cause of cold [59]. Similarly, it has been well documented that the majority of HCoV-NL63 infections are mild in humans, though occasionally this coronavirus causes pneumonia or central nervous system diseases in susceptible individuals [60]. During 2009 and 2016, it accounted for about 0.5% of all acute respiratory tract infections in hospitalized patients from Guangzhou, China, but few death cases are reported [61]. HCoV-229E is a close relative of HCoV-NL63 and it will lead to alike symptoms [62].

Figure 3 displays the CGR plots of different sequences of human coronavirus at the value of 6 for k-mer frequency. The CGR plots visually indicate that the genomic signature of the SARS-CoV-2 isolate Wuhan-Hu-1 (Fig. 3c) is closer to the genomic signature of the SARS-CoV coronavirus isolate Canada (Fig. 3a), followed by the strain of MERS-CoV Betacoronavirus England 1 isolate (Fig. 3b). Moreover, the other four human coronaviruses from Fig. 3d (HCoV-OC43/JN129835.1), Fig. 3e (HCoV-HKU1/NC_006577.2), Fig. 3f (HCoV-229E/JX503060.1) and Fig.3g (HCoV-NL63/JQ765575.1) presents similar visual patterns, which are different from the former three types. Given the CGR plots of human coronaviruses, we further explore the trace of their origin and relation through phylogenetic analysis. We randomly select five complete genomes from each type containing the reference strain. The phylogenetic tree is constructed based on all pairwise distance with maximum likelihood technique for the dataset. The results in Fig. 4 presents a clear separation of seven clusters and relationships within the clusters. The average inter-cluster distances confirm that SARS-CoV-2 sequences are closest to the species of SARS-CoV (average distance 0.486), followed by MERS-CoV (4.782), which are far away from other four human coronaviruses. We also find that HCoV-OC43 and HCoV-HKU1, HCoV-229E and HCoV-NL63 may originate from the same ancestor with the genetic distance 1.842 and 2.779, respectively. But there is no evidence indicating the situation that the two different species of human coronavirus will present similar lethality if they are genetically close.

**Figure 3.**
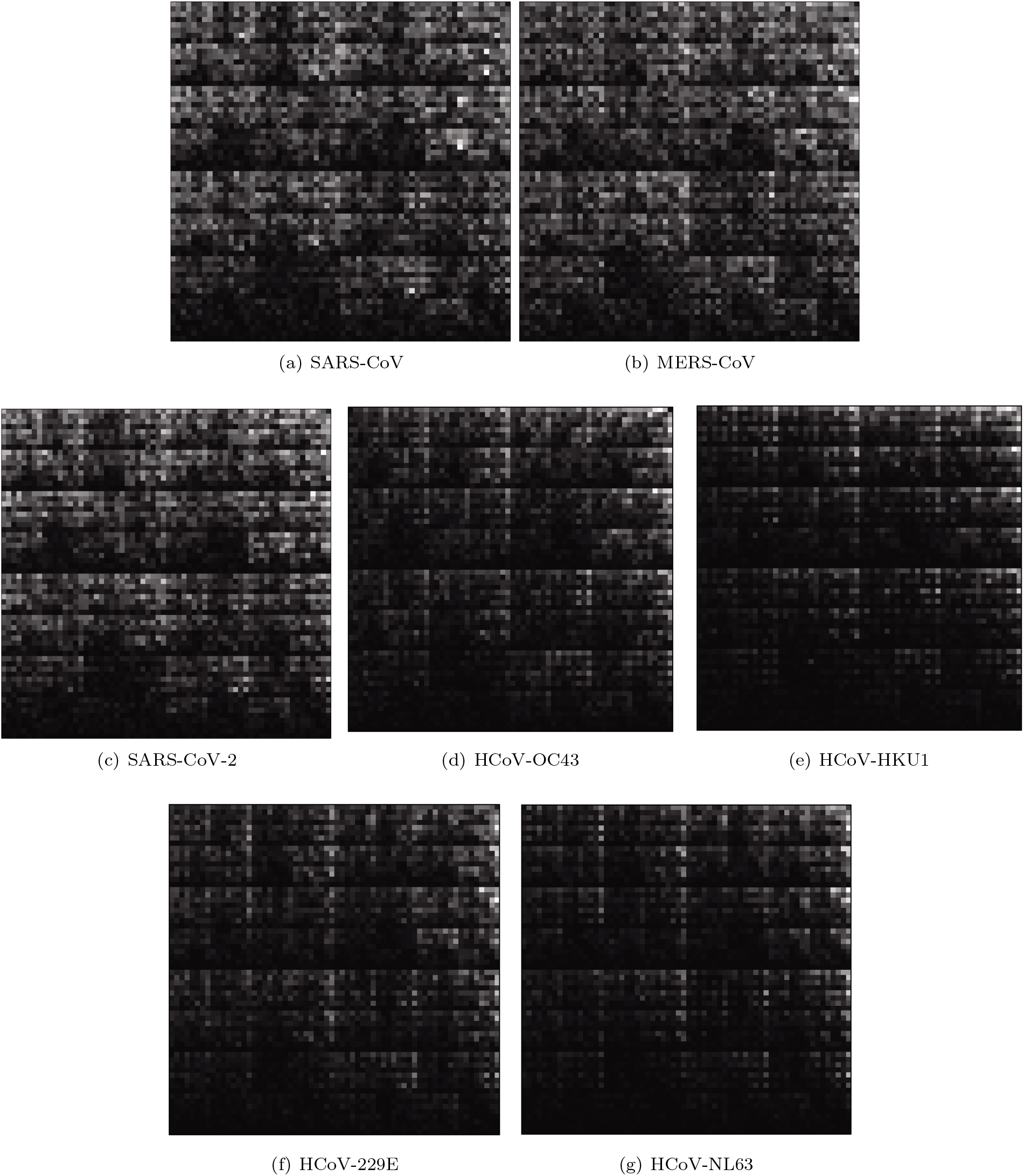
The visualization of Chaos Game Representation plots of references of seven human coronavirus at *k* = 6 of (a) SARS-CoV/NC_004718.3/Severe acute respiratory syndrome-related coronavirus/Canada, (b) MERS-CoV/NC_038294.1/Betacoronavirus England 1, Middle East respiratory syndrome-related coronavirus/United Kingdom, (c) SARS-CoV-2/NC_045512.2/Severe acute respiratory syndrome coronavirus 2 isolate Wuhan-Hu-1/China, (d) HCoV-OC43/JN129835.1/Human coronavirus OC43 strain HK04-02, China/Betacoronavirus 1, (e) HCoV-HKU1/NC_006577.2/Human coronavirus HKU1, (f) HCoV-229E/JX503060.1/Human coronavirus 229E isolate 0349, Netherlands, (g) HCoV-NL63/JQ765575.1/Human coronavirus NL63 strain NL63/DEN/2005/1876, USA. The vertices of the plot are assigned A (top left), T (top right), C (bottom left), G (bottom right).

**Figure 4.**
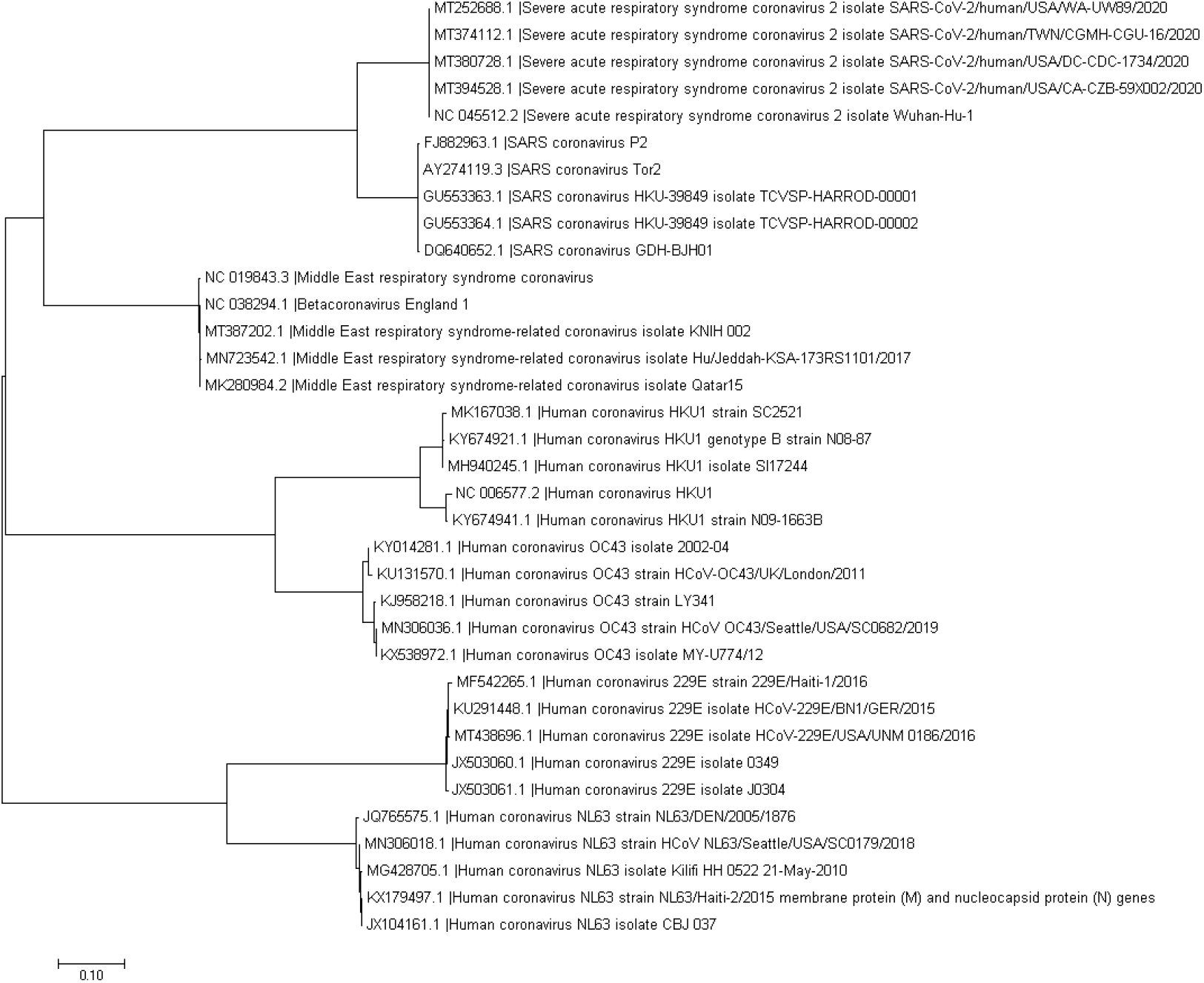
The phylogenetic tree using maximum likelihood generated pairwise distance matrix shows the hierarchical classification between different human coronaviruses.

## Discussion

We provided a comprehensive quantitative analysis to predict the lethality of potential new human-adapted coronavirus with six different numerical representations of RNA sequences applied in machine learning models. The models are computational efficiency because they are alignment-free. Compared with alignment-based methods, multiple sequence alignment is not needed with the leverage of DFT techniques. The experiment results show that most of the models have achieved rapid and accurate predictions for the lethality of new human-adapted coronavirus. We validated our results by a quantitative analysis based on the construction of the phylogenetic tree, which reveals the evolutionary relationships among all human coronaviruses based upon genetic information. Coronaviruses are usually thought to cause mild and non-lethal symptoms in humans before the outbreak of SARS-CoV in 2003. The high pathogenicity of SARS-CoV, MERS-CoV and newly SARS-CoV-2 captures surgent interests and concerns of the family of coronavirus. Timely analysis of genomic sequences of novel strains requires quick sequence similarity comparison with thousands of known species, which are generally performed by alignment-based methods. However, these methods are time-consuming and sometimes challenging in cases where homologous sequence continuity cannot be ensured. The application of alignment-free approaches has addressed this issue that can handle a large number of sequences effectively.

Previous studies have elucidated that the origin of this SARS-CoV-2 stems from bats [10] [63]. Early sequencing of SARS-CoV-2 strains revealed over 99% similarity with some bat-like coronavirus, indicating these infections result from a recent cross-species event [64]. Bats are regarded as the natural reservoir of viruses and cross-species transmission to mammals [4] [65]. Before the emergence of SARS-CoV-2, it is uncovered that the coronavirus SARS-Cov and MERS-CoV have also originated from bats [66] [67]. The phylogenetic analyses assist to identify the relationships between SARS-CoV-2 and other coronaviruses through nucleotide and amino acid sequence similarities. The continuous human-to-human transmission has been confirmed and asymptomatic cases have continued to increase [68] [69]. There is a desperate need for strict precautions to prevent the spread of the virus and protect public health. Vaccines and miracle drugs are the most efficient ways of fighting against this crisis. Currently, the development of vaccines has been into Phase 3 trials in some countries, while the human ACE2 receptor has been identified as the potential receptor for COVID-19 and serves as a potential target for treatment [70] [71]. Nevertheless, with the circulation of bat-related coronavirus and geographic coverage, it is critical to monitor the evolution of coronavirus. Currently, seven known types of coronavirus can infect humans. Novel strains of these coronaviruses can likely arise and attack human again through reassortment and mutation when two different or more strains co-infect the same host. Preparation is necessary to prevent potential epidemics and pandemics caused by a novel coronavirus. As a result, our work paves the basis for surveillance by inferring the lethality of any potential human coronaviruses that may emerge in the future.

This study is subject to a variety of limitations. The definition of classifying the degree of coronavirus lethality is mainly based on the mortality rate. We assume that the higher the mortality, the more lethal for the virus, and thus make three categories of the lethality level for all viruses with a different threshold. However, our estimation for these values lies within the range of fatality rate from the literature, which we do not have sufficient data to parameterize the case-structured model, especially for viruses with few samples. We also do not build a benchmark for the death caused directly by human coronaviruses, as the criteria from institutions and countries could be different. Besides, the limited data points for the human coronavirus pale the high predictive accuracy, as most of the machine learning algorithms possess a superb generation ability to discover inherent patterns from training samples, particularly in the small dataset. But like typical machine learning approaches, our models are not qualified to provide a direct and accessible explanation that explicitly interprets why a certain coronavirus strain is more lethal to humans. Some rule-based methods or clinical study might provide a better rationale for their results.

## Conclusion

We provide a comprehensive analysis through alignment-free machine learning-based methods for the prediction of the lethality of potential human-adapted coronavirus. The results show that on the average, CGR, EIIP, and Just-A representations perform better than others. Interestingly, traditional machine learning methods display obvious merit both in computational efficiency and performance than deep learning models on this task. Validation of other types of human coronavirus in combination with phylogenetic analysis further demonstrates our predictive results. We hope this work would facilitate the research of COVID-19 for biologists and clinicians that are in the frontline.

## Acknowledgments

This project is supported by AcRF Tier 2 grant MOE2014-T2-2-023, Ministry of Education, Singapore and A*STAR-NTU-SUTD AI Partnership grant, RGANS1905.

